# Hybridisation boosts dispersal of two contrasted ecotypes in a grass species

**DOI:** 10.1101/2021.04.16.440116

**Authors:** Emma V. Curran, Matilda S. Scott, Jill K. Olofsson, Florence Nyirenda, Graciela Sotelo, Matheus E. Bianconi, Sophie Manzi, Guillaume Besnard, Pascal-Antoine Christin

## Abstract

- In the absence of strong reproductive barriers, genetic exchanges between closely related groups of organisms with different adaptations have well-documented beneficial and detrimental consequences. In plants, pollen-mediated exchanges affect the sorting of alleles across physical landscapes, and influence rates of hybridisation. How these dynamics affect the emergence and spread of novel ecological strategies remains only partially understood.
- We use phylogenomics and population genomics to retrace the origin of two geographically overlapping ecotypes of the African grass *Alloteropsis angusta*.
- We report the existence of a previously undescribed ecotype inhabiting miombo woodlands and grasslands. The two ecotypes have divergent nuclear genomes. However, the seed-transported chloroplast genomes are consistently shared by distinct ecotypes inhabiting the same region. These patterns suggest that the nuclear genome of one ecotype can reach the seeds of the other via pollen movements, with strong selection subsequently sorting nuclear alleles by habitat.
- The contrasting ecotypes of *A. angusta* can use each other as a gateway to new locations across a large part of Africa. Coupled with newly discovered hybridisation with the sister species *A. semialata*, our results show that hybridisation can facilitate the geographical dispersal of distinct ecotypes of the same grass species.

## Introduction

When populations encounter new environments, the selection pressures they face may initiate the evolution of novel adaptive phenotypes, enabling them to colonise previously untapped niches. Contrasting conditions between novel and ancestral habitats can impose divergent selection on populations, resulting in reproductively isolated ecotypes within a species (Schluter, 2001). Such ecotypes may represent early stages of the ‘speciation continuum’ (Seehausen *et al*., 2014), and, given time, they may become fully isolated species. Prior to complete reproductive isolation, however, hybridisation between diverged ecotypes is commonplace (e.g. in monkeyflowers: Stankowski *et al*., 2017; marine snails: Galindo *et al*., 2013; and whitefish: Lu *et al*., 2001). Hybridisation between geographically isolated populations may result in a breakdown of differentiation, leading to loss of local adaptation and the formation of homogenised ‘hybrid swarms’ (Taylor *et al*., 2006; Abbott *et al*., 2013). Hybridisation can also contribute to the overall success of populations. Adaptive genetic material may be shared between populations via introgression, conferring advantageous traits (e.g. Song *et al*., 2011; The Heliconius Genome Consortium, 2012; Arnold *et al*., 2016). In sessile organisms, such as plants, hybridisation can affect the spatial movements of genes (Martinsen *et al*., 2001; Petit *et al*., 2003). Plant pollen can be exchanged over long distances, resulting in hybridisation with both positive and negative consequences (Ellstrand, 1992, 2014). Indeed, long-distance cross pollination can restore diversity, but also lead to the swamping and extinction of rare types (Ellstrand, 1992; Aguilée, *et al*., 2016; Hall, 2016; Todesco *et al*., 2016). However, the dynamics that govern barriers to gene flow within plant species, i.e. the balance between gene flow and divergent selection (Barton & Hewitt, 1985), and the consequences of recurrent hybridisation are still not fully understood.

Grasses occur in most ecosystems around the world, a feat facilitated by the flexibility of their growth plan and variation in life strategies (Linder *et al*., 2018). They range from short annuals to large, tree-like bamboos, and include mat-forming alpine species, aquatic plants, and decumbent species crawling over large areas (Grass Phylogeny Working Group, 2001; Perreta *et al*., 2011; Kellogg, 2015). However, the evolutionary dynamics underlying the origins of different growth forms remain poorly studied. The grass genus *Alloteropsis* is composed of five recognised species (Clayton and Renvoize 1982; Ibrahim *et al*., 2008). The two sister species *A. semialata* and *A. angusta* are perennials. The former inhabits the grasslands and savannah woodlands of Africa, Asia and Oceania (Lundgren *et al*., 2015), and its stems grow erect from bulb-like structures formed by thickened sheaths (Clayton and Renvoize 1982). The plant propagates vegetatively via very short rhizomes that lead to a proliferation of bulbs next to each other. By contrast, *A. angusta* is reported as a slender, decumbent species growing in swamps, and is only known from Central and East Africa (Stapf 1919; Clayton and Renvoize 1982). While studying the origins of novel photosynthetic types in *A. semialata*, we discovered some specimens assigned to this species that, based on both organelle and nuclear genomes, corresponded to *A. angusta* (e.g. sample “Pauwels 1182 [BRU]”; Olofsson *et al*., 2016). These misidentified samples were erect, with bulb-like enlargements at the base of the stem, showing that *A. angusta* can occur as solid erect plants that resemble *A. semialata* in addition to the previously reported fragile decumbent type. Because the growth forms observed in the field persisted when plants were grown in controlled conditions at the University of Sheffield, they must be genetically determined. *Alloteropsis angusta* therefore constitutes an outstanding system to study the evolutionary dynamics leading to habitat specialisation of the growth form within grass species. Since this species has not yet been the focus of dedicated studies, the status of the two morphs and their relationship remain unknown.

In this study, we combine phylogenomics and populations genomics to study the dynamics underlying the functional diversification of *A. angusta* in Africa. We quantify the morphological variation within *A. angusta* to (i) confirm the existence of two morphs associated with different environments. We then sequence the genomes of 13 individuals of *A. angusta* and infer the phylogenetic tree of chloroplast genomes to (ii) determine whether maternally-inherited genomes of each morph spread independently. Phylogenetic analyses of the nuclear genomes from the same individuals are then used to (iii) infer the relationships among the two morphs. We scan the genomes of numerous populations spread from Zambia to Uganda and use population genomics to (iv) test for frequent admixture between the two types and assess the geographical patterns of genetic variation. Finally, ABBA-BABA tests are conducted to (v) test for introgression between the two morphs and with the sister species *A. semialata*. Our results support the importance of frequent hybridisation among the two morphs of *A. angusta* for the dispersal of this species, despite a strong ecological barrier. We also reveal episodic exchanges between *A. semialata* and one of the morphs of *A. angusta*, suggesting that the two species might form a loose species complex.

## Materials and Methods

### Population sampling

Samples of *Alloteropsis angusta* were obtained from herbaria, or from fieldwork conducted in Uganda, Tanzania and Zambia (Table S1; Olofsson *et al*., 2016; Dunning *et al*., 2017). Denser, population-level sampling was conducted in Zambia through walk-and-search stops in areas covered by miombo woodlands, grasslands or swamps (Table S1). For each population found, GPS coordinates were recorded with a description of the habitat, and up to ten distinct individuals, growing at least one metre apart, were collected in silica gel. For most populations, several individuals were pressed and later used to prepare herbarium vouchers (listed in Table S1). When *A. angusta* and *A. semialata* grew together, both were sampled. In addition, we sampled 46 plants with GPS coordinates for each individual in a location where contrasted morphs were observed (population ZAM1930).

### Morphological analyses

The 80 available herbarium vouchers were digitised and measured with ImageJ (Schneider *et al*., 2012; Table S2). To capture variation in vegetative organs, we measured the size of the bulb, stems and leaves from all samples. The bulb length was measured from its bottom extremity to the limit between protective sheaths and rosette leaves, while the bulb width was measured as the largest width perpendicular to the growth axis. The stem length was measured from the limit of the bulb to the split of the racemes, and its width was measured at the base, just above the bulb, and at the top, just below the racemes. For leaves, the length of the lamina of the longest rosette leaf and longest stem leaf were measured. To capture reproductive characters, the length of the longest raceme was measured, together with the average length of ten spikelets and the average spacing between all consecutive spikelets along a raceme. The anatomical variation was summarised with a principal component analysis (PCA), considering all characters except the spikelet length, which could not be typed on individuals that lost their florets before collection. The PCA was performed using the *prcomp* function in R version 3.6.0 (R Core Team, 2019).

### Genome sequencing and phylogenetic analysis of chloroplast genomes

Phylogenetic relationships within and between the two sister species were inferred using whole genome re-sequencing data. The genomes of two herbarium samples and three field-collected samples were sequenced using a genome skimming approach, with 150-bp paired-end Illumina reads, as described in Olofsson *et al*. (2016). In addition, four field-collected individuals representing a diversity of geographic origins and growth habits were sequenced as 250-bp paired-end reads, as described in Dunning *et al*. (2019) (Table S1). Other genome datasets of *A. angusta, A. semialata* and *A. cimicina* were retrieved from previous work (Table S1).

Complete chloroplast genomes were assembled using the genome walking approach described in Lundgren *et al*., 2015. Assembled plastomes were aligned using ClustalW, the second repeat was removed, and the alignment was manually refined. The final alignment was 120,271 bp long, and a time-calibrated phylogenetic was inferred with Beast v. 1.8.4 (Drummond and Rambaut, 2007). A log-normal relaxed clock was used, with a GTR+G substitution model and a constant-size coalescent prior. The monophyly of the ingroup (all samples other than the outgroup *A. cimcina*) was enforced to root the tree. The root of the tree was set to 11.46 Ma and the split of *A. angusta* and *A. semialata* to 8.075 (using a normal distribution of 0.0001), following previous estimates (Lundgren *et al*., 2015). Two analyses were run for 50,000,000 generations, sampling a tree every 10,000 generations. Convergence of the runs was monitored using Tracer v. 1.6.0 (Rambaut *et al*., 2013), and the burn-in period was set to 10,000,000 generations. The median ages of posterior trees were mapped on the maximum credibility tree.

### Phylogenetic analyses of nuclear genomes

Nuclear variants were extracted from sequencing reads of whole-genome sequenced samples. Raw reads obtained from fresh samples were cleaned using NGS QC Toolkit v. 2.3.3 (Patel & Jain, 2012) to remove reads where more than 20% of the bases had a quality score below Q20, and those which had ambiguous bases. Bases with quality score below Q20 were further trimmed from the 3’ end of reads. Adaptors were removed using NxTrim (O’Connell *et al*., 2015). Raw reads from low coverage sequenced individuals were cleaned as in Olofsson *et al*. (2016). All cleaned reads were aligned to the *A. semialata* reference genome (ASEM_AUS1_v1.0; GenBank accession QPGU01000000; Dunning *et al*., 2019) using Bowtie2 v. 2.3.5.1 (Langmead & Salzberg, 2012) with the default settings for paired-end reads. Read alignment files were cleaned, sorted and indexed using samtools v.1.9 (Li, 2011). PCR duplicates were removed using Picard Tools v.1.102 (Broad Institute, 2019).

For each sample, reads mapped to the reference nuclear genome were separated out from those mapped to the reference mitochondrial or chloroplast genomes. A multi-sample sequence alignment was then generated across all individuals. A consensus sequence was generated from the reference-mapped reads for each individual using the mpileup tool in samtools v.1.9 (Li, 2011), and a custom bash script was then used to filter for depth the consensus sequences of re-sequenced individuals (Olofsson *et al*., 2016), removing bases supported by less than three reads. The genome-skimmed individuals were not filtered for depth, due to the low average coverage for these samples. Polymorphic sites were called as ambiguous bases following IUPAC codes. Sites with more than 10% missing data were removed using trimAl v. 1.4.rev6 (Capella-Gutiérrez *et al*., 2009), resulting in an alignment of 142,848 bp. A maximum likelihood phylogeny was estimated under the GTR+CAT substitution model in RAxML v. 8.2.10 (Stamatakis, 2014) with 100 bootstrap pseudoreplicates.

We also estimated a multigene coalescent phylogeny from the nuclear genomes. For this, we first retrieved a dataset consisting of 7,408 putative single-copy orthologs of Panicoideae grasses (the tribe including *Alloteropsis*) from a previous study (Bianconi *et al*., 2020), and the corresponding coding sequences of *A. semialata* were used here as references for the assembly of gene sequences, as described for the whole genome. The resulting gene alignments were trimmed using trimAl to remove sites with more than 30% missing data, and sequences shorter than 200 bp after trimming were discarded. Finally, only gene alignments longer than 500 bp and with more than 95% of taxon occupancy were retained for phylogenetic analyses. A maximum-likelihood phylogeny was estimated for each of the remaining 2,960 alignments using RAxML, with a GTR+CAT substitution model and 100 bootstrap pseudoreplicates. Gene trees were then summarised into a multigene coalescent phylogeny using Astral v5.5.9 (Zhang *et al*., 2018) after collapsing branches with bootstrap support below 50.

### *Population-level genetic structure within* A. angusta

The genomes of population-level samples were scanned using a reduced representation approach, restriction site associated DNA (RAD) sequencing, as described in Olofsson *et al*. (2019). In brief, the DNA of up to five individuals of each population (more in ZAM1930 where the two morphs occurred) were double digested and then pooled before sequencing 72 – 96 individuals per lane of Illumina HiSeq 2500 at the Sheffield Diagnostic Genetics Service, Sheffield Children’s NHS Foundation Trust. Our final dataset included 196 *A. angusta* individuals, and 15 newly sequenced *A. semialata*, which were combined with 49 previously sequenced *A. semialata* (Olofsson *et al*., 2021; Table S1).

We used a PCA of genetic variation and individual ancestry analyses to describe the distribution of genetic diversity among these populations. The program Trimmomatic (Bolger *et al*., 2014) was used to trim raw RAD-sequencing reads to remove adaptor and other Illumina-specific sequences and bases with a low quality score (Q < 3) from the 5’ and 3’ ends. Reads were further clipped when the average quality within a sliding window of four bases dropped below an average quality threshold (Q < 15). The pooled reads were de-multiplexed into individually barcoded samples using the module ‘process_radtags’ in the program STACKS (Catchen *et al*., 2013). Reads were then mapped to the *A. semialata* reference genome (ASEM_AUS1_v1.0; GenBank accession QPGU01000000; Dunning *et al*., 2019), using Bowtie2 with the default settings for paired-end reads. We estimated genotype likelihoods from the reference-mapped reads for each individual, using the program ANGSD (Korneliussen *et al*., 2014). Sites present in at least 50% of the individuals with a minimum depth of 5 per individual and with minimum mapping and base quality scores of 20 were included in the analysis. Individuals with more than 99% missing data were removed. After filtering, 49,523 sites remained, across 196 individuals. The proportion of each individual’s genome that can be assigned to a specified number of genetic clusters (*K*) was estimated from the genotype likelihoods using the software NGSadmix (Skotte *et al*., 2013). NGSadmix was run with *K* ranging from 1 to 10, with five replicates for each run, each with a random starting seed. The explanatory power of the increasing *K* values was assessed using the Δ*K* criterion (Evanno *et al*., 2005), implemented with the CLUMPAK tool (Kopelman *et al*., 2015). A PCA was carried out using PCAngsd (Meisner & Albrechtsen, 2018), which estimates a covariance matrix using the genotype likelihoods. We retrieved the principal components of genetic structure using eigenvector decomposition in R.

To establish the genetic structure within and between the erect and decumbent morphs of *A. angusta*, we calculated pairwise genome-wide *F*_ST_ between all *A. angusta* populations using Hudson’s *F*_ST_ estimator (Hudson *et al*., 1992). This was calculated for each SNP based on allele frequencies estimated directly from the data, as in Soria-Carrasco *et al*. (2014). Average genome-wide *F*_ST_ was calculated as a ratio of averages, by averaging the within- and between-population variance components separately as recommended by Bhatia *et al*. (2013), using an R script by Soria-Carrasco (2019: https://github.com/visoca/popgenomworkshop-hmm). The erect and decumbent individuals of population ZAM1930 were considered as two distinct populations for these analyses. Pairwise geographic distances between pairs of populations were calculated using the ‘rdist.earth’ function in the R package ‘fields’ (Nychka *et al*., 2017). A relationship between genetic and geographic distances was tested using Mantel tests, with 9,999 permutations, for populations within each of the erect and decumbent groups, and among them. In each case, the strength of the relationship was assessed with Spearman’s correlation coefficient (ρ).

To examine the genome-wide landscape of genetic differentiation between the decumbent and erect morphs, we used the *F*_ST_ calculated for each SNP between them in population ZAM1930, for which there are 20 decumbent and 26 erect individuals. Average *F*_ST_ was calculated, as above, for non-overlapping sliding windows of 500kb.

### *Tests for gene flow between morphological types of* A. angusta *and between species*

To test for gene flow among *A. angusta* lineages and between *A. angusta* and *A. semialata*, we used the ABBA-BABA method (Durand *et al*., 2011; Green *et al*., 2010). This approach uses patterns of ancestral and derived alleles to calculate *D*-statistics and test for asymmetry in the frequencies of phylogenies incongruent with the species tree beyond what is expected under a scenario of incomplete lineage sorting (Durand *et al*., 2011; Green *et al*., 2010). Three ingroups (P1, P2, P3) and one outgroup (O) are required, in the configuration (((P1, P2), P3), O), and here *A. cimicina* was used as the outgroup. Tests were carried out using the –doAbbababa option in the program ANGSD (Korneliussen *et al*., 2014), to compute the *D* statistic. Deviation from the null expectation (D = 0) was tested using the jackKnife.R script (block jackknife method) provided with ANGSD.

To test for gene flow among *A. angusta* lineages, we considered first all combinations where P1 and P2 are occupied by a decumbent individual and P3 by an erect individual, and then all combinations where P1 and P2 are occupied by an erect individual and P3 is occupied by a decumbent individual. The Bonferroni correction was used to adjust p-values with the total number of such combinations, but only a subset was subsequently considered to simplify the dataset. For each of the two types of combinations, the individual most often found as the least introgressed was identified and then fixed in the P1 position, so that all positive *D*-statistics indicate gene flow between P2 and P3. A similar approach was used to test for gene flow between *A. angusta* and *A. semialata*, considering all combinations where P1 and P2 are occupied by *A. angusta* individuals and P3 is an individual from *A. semialata*. Again, p-values were corrected for the number of such comparisons, but only those with the least often introgressed individual in the P1 position were considered.

## Results

### Two growth forms associated with distinct habitats

We collected six populations of *A. angusta* in 2015 in Uganda, in boggy areas around lakes or other water soaked grasslands. All these individuals were decumbent, with long fragile stems crawling among other grasses. The stems were branching, with secondary roots growing from nodes (e.g. Fig. 1C). Individuals brought back as live cuttings have been living in controlled conditions for > six years, with the same growth habit. They spread horizontally by making new roots from crawling stems that can then produce new individuals. Using the approach described in Bianconi et al. (2020), we estimated the genome size for two decumbent individuals, and the estimated values were similar to diploids of the sister species *A. semialata* (individual MRL15-04-06, 2C = 2.09 Gb; MRL15-04-08, 2C = 1.95 Gb). The individuals of *A. angusta* collected in Tanzania in 2016 and Zambia in 2017 were clearly distinct from the Ugandan ones. They grew in miombo woodlands and associated grasslands from small bulbs, each with a single erect stem (e.g. Fig. 1B). We grew one individual from seed in controlled conditions (from population ZAM1720), and it produced a bulb and grew erect, as in the field. Its genome size was similar to that of Ugandan individuals (2C = 2.10 Gb).

**Figure 1.**
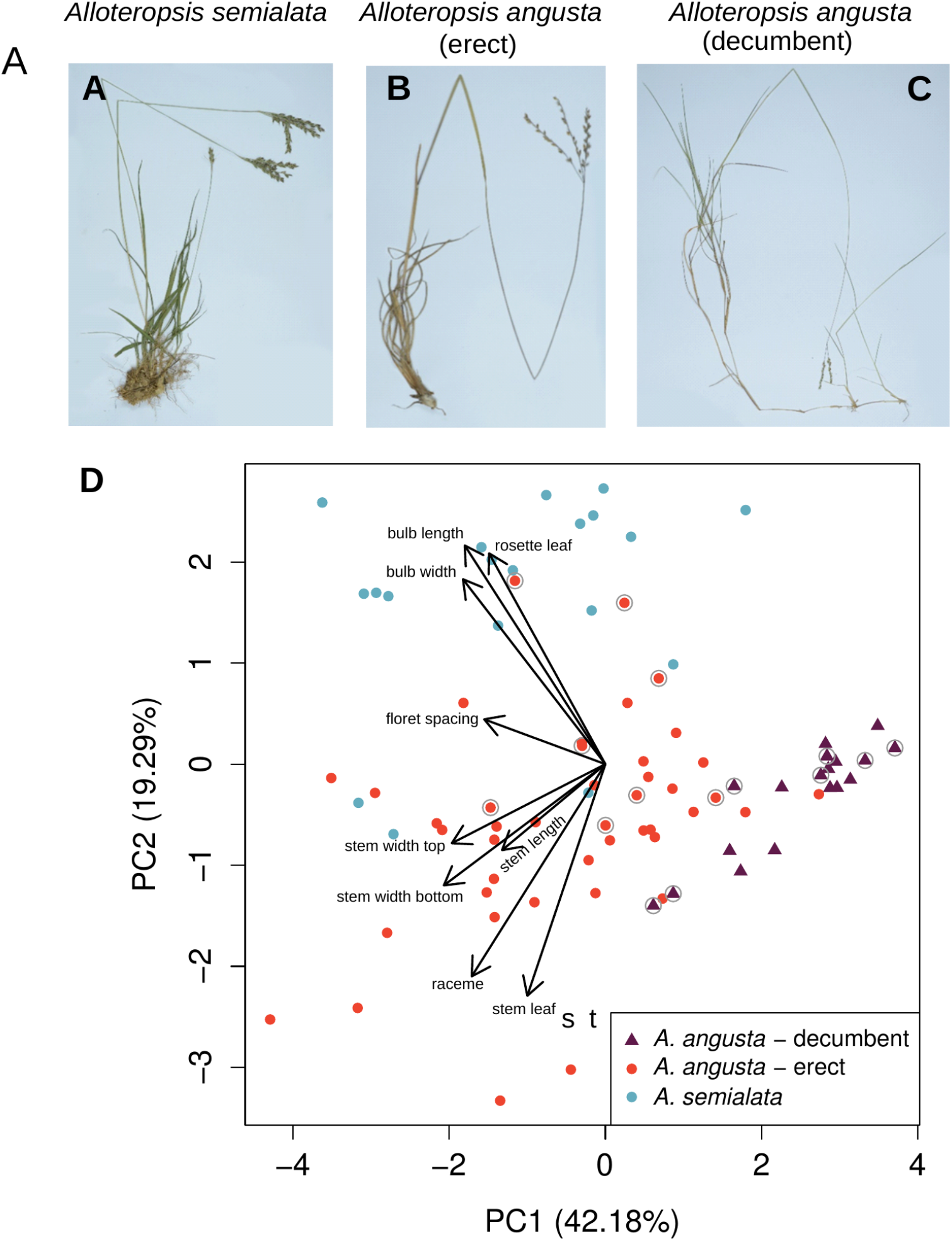
Morphological variation within *Alloteropsis angusta* and its sister species *A. semialata*. Digitised herbarium specimens of **A)** *A. semialata* [ZAM1503 – Nyirenda, Lundgren Dunning 3 (SHD)], **B)** *A. angusta –* erect ecotype [TAN1601 – Dunning, Dunning, Kayombo 1 (SHD)], and **C)** *A. angusta* – decumbent ecotype [M. R. Lundgren 2015-3-3 (SHD)]. **D)** Morphological variation among *A. semialata* and *A. angusta*, as assessed by the first two axes of a principal component analysis. Contributions of the different variables are indicated with arrows. Individuals with a grey circle are included in the whole-genome phylogeny.

In total, we collected 31 populations of *A. angusta* in Zambia and Tanzania. In four of these populations, the individuals were described as decumbent in the field, with strongly branching stems, crawling amongst other species. All of these decumbent individuals, which were morphologically similar to those collected previously in Uganda, were found in water-logged wetlands on the shores of rivers or lakes (Table S1; Fig. S1). Individuals from 26 of the other populations were described in the field as erect. They formed bulb-like structures of various sizes at the base of their stems and grew tall, with a single stem per bulb. Short rhizomes connected underground successive bulbs. These individuals match the description of the defunct species *A. gwebiensis* (Stent & Rattray, 1933), which was later merged with *A. semialata* (Clayton & Renvoize, 1982). These erect individuals were all found in miombo woodlands and associated grasslands, with between 0 and 90% tree cover (Table S1; Fig. S1). While two of these erect populations were found in grasslands topping rivers, these grasslands were not directly connected to the rivers and did not present the boggy characteristics of the sites in which the decumbent forms were found. In seven of the erect populations, *A. angusta* grew together with the conspecific *A. semialata* (Table S1). Finally, population ZAM1930 spanned a miombo woodland sloping toward a river wetland. Individuals growing in the miombo of this site were erect and found mixed with *A. semialata*. By contrast, the individuals growing in the open river wetland were all decumbent (Fig. S2).

### The growth forms are morphologically distinct

The morphology of all sampled *A. angusta*, including herbarium vouchers, was analysed together with a number of *A. semialata* (Table S2). The spikelet length of *A. semialata* ranged from 4.28 to 6.32 mm, while that of the decumbent *A. angusta* ranged from 2.7 to 3.75 mm (Table S2). These values correspond to the previously described *A. semialata* and *A. angusta* (Clayton & Renovize 1982), but the spikelet length of the erect *A. angusta* overlapped with the two groups (3.13-4.52 mm; Table S2). A PCA confirmed that the erect and decumbent forms of *A. angusta* occupy different parts of the morphological space (Fig. 1D), although the growth habit was not included as a character. Important variation was observed within the erect *A. angusta*, which overlap in the anatomical space with *A. semialata* (Fig. 1D). Populations classified as decumbent occurred in a small subset of the anatomical space characterised by shorter and thinner bulbs, smaller rosette leaves, and thinner stems (Fig. 1D). The bulb width clearly discriminated the two types, being 0.90-2.57 mm in the decumbent *A. angusta* and 3.06-12.81 mm in the erect ones (5.88-19.13 mm in *A. semialata*; Table S2). The presence of bulbs is therefore a suitable character to define the two growth habits of *A. angusta*, which are consistently associated with contrasted habitats and therefore correspond to ecotypes (Figs 1, S1).

### Chloroplast genomes are shared by the two ecotypes

The individuals selected for whole genome sequencing capture the morphological diversity within the group (Fig. 1). The chloroplast phylogeny sorts *A. angusta* accessions by geographic origin, independently of their ecotype (Figs 2B, S3). The first split separates two groups, each with erect and decumbent individuals, and the decumbent accessions form four distinct clades, while the erect accessions form five (Fig. 2B). In particular, the erect and decumbent individuals from the West of Zambia (ZAM2074-14 and ZAM2075-04) are grouped together (Fig. 2B). The two contrasted types from population ZAM1930 (ZAM1930-JKO0102 and ZAM1930-17) are also grouped, with almost no chloroplast divergence (one substitution, four 1-bp indels, and one 19-bp indel out of 117,652-bp pairwise alignment). These patterns indicate that the history of the maternally-inherited chloroplasts is shared between the erect and decumbent ecotypes. The divergence times among the plastomes of *A. angusta* individuals are proportional to geographical distances (Mantel test, *ρ* = 0.67, *P* < 0.001), indicating that the extant plastome diversity results from a gradual expansion. The relationship remains significant when considering only pairs composed of the two ecotypes (Mantel test, *ρ* = 0.57, *P* < 0.001), and the patterns are similar for the different types (Fig. S4).

**Figure 2.**
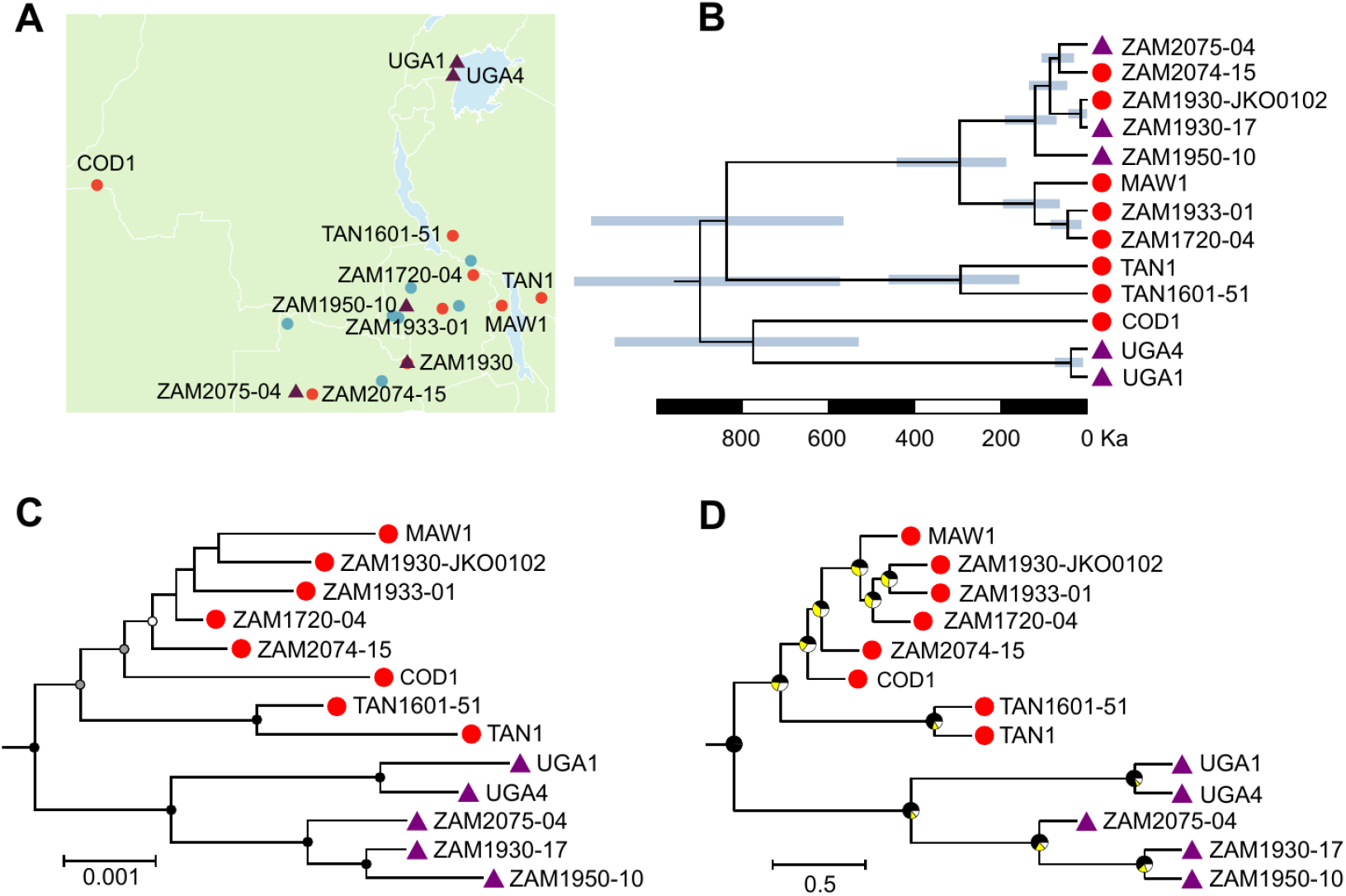
Phylogenetic relationships among *Alloteropsis angusta* samples. **A)** The geographic distribution of individuals of *Alloteropsis angusta* and *A. semialata* included in the phylogenetic tree is shown. Erect individuals of *A. angusta* are shown with red circles, and decumbent individuals with purple triangles. Samples of *A. semialata* are shown with blue circles. **B)** The portion of the time-calibrated phylogenetic tree corresponding to *A. angusta* is shown (see Fig. S3 for full tree). Bars at nodes indicate 95 %-confidence intervals. All nodes had support values of 1.0. The scale is given in thousands of years (Ka). **C)** The portion of the maximum-likelihood tree based on nuclear variants corresponding to *A. angusta* is shown (see Fig. S3 for full tree). Dots on nodes show bootstrap support values; 100 = black, > 90 = grey, > 60 = white. **D)** The portion of the multigene coalescence species tree corresponding to *A. angusta* is shown (see Fig. S4 for full tree). Pie charts on nodes show the proportions of quartets supporting the main topology (in black) and the two alternatives. All nodes have support values above 0.97. The scale is given in coalescence units. Terminal branches are given arbitrary lengths. The scale is given in expected substitutions per site. For **B), C)**, and **D)**, ecotypes are shown at tips; red circles = erect individuals, purple triangles = decumbent individuals.

### Deep nuclear divergence of the two ecotypes

In stark contrast to the chloroplast phylogeny, the maximum likelihood nuclear phylogeny using whole genome sequenced individuals revealed that the erect and decumbent ecotypes each form a distinct monophyletic group that covers the studied geographical region (Figs 2C, S3). Individuals of contrasting ecotypes that were collected from the same locality in Zambia (population ZAM1930) group with their respective ecotype despite close geographic proximity, with the decumbent individual (ZAM1930-17) showing a closer phylogenetic relationship to decumbent individuals from Uganda (UGA1 and UGA4) than to erect individuals growing a few metres away (ZAM1930-JKO0102).

The multigene coalescent species tree recovered the same relationships (Figs 2D, S5). The monophyly of *A. angusta* was supported by almost all tree quartets, indicating lineage sorting on the branch leading to *A. angusta*. By contrast, many quartet trees supported the two topologies alternative to the main one for nodes leading to the erect *A. angusta* (Fig. 2D), which indicates that high levels of incomplete lineage sorting occurred within *A. angusta*. The amount of quartets supporting alternative topologies was much smaller in nodes of the decumbent group (Fig. 2D). Overall, these phylogenetic patterns suggest that the nuclear genomes of the two ecotypes diverged before the spread of *A. angusta* across Africa.

### Nuclear genetic groups are maintained despite close geographic proximity

The genetic structure of *A. angusta* populations spread across Zambia, Tanzania and Uganda was deciphered using population-level RAD-sequencing data (Fig. 3A). The largest component of genetic variation separates the erect and decumbent ecotypes along the first axis (23.9% of the total variation, Fig. 3B). The second axis (9.71% of the variation) splits decumbent individuals according to geography, separating the more distant Ugandan populations from the rest of the individuals (Fig. 3B). The number of genetic clusters best describing the population structure within *A. angusta* is two (Fig. S6), again sorting the samples according to ecotype (Fig. 3C). Multiple decumbent individuals show suggestions of introgression with the erect group (Fig. 3C), consistently across individuals of the decumbent population ZAM2075 and the erect populations ZAM2074, ZAM2095, and ZAM2069. Overall, across populations spread from Uganda to Zambia, the two ecotypes behave as distinct nuclear groups (Fig. 3C). The structure remains strong between the ecotypes even within the population where they grow adjacent to one another (ZAM19-30; Figs 3C, S2), and again decumbent samples from this population are assigned to the same genetic cluster as Ugandan samples growing >1400 km away, instead of erect individuals growing a few meters away. The nuclear patterns observed across the species range (Figs 2C, 2D) therefore translate to smaller scales, with geographically close ecotypes associated with distinct and divergent nuclear genomes.

**Figure 3.**
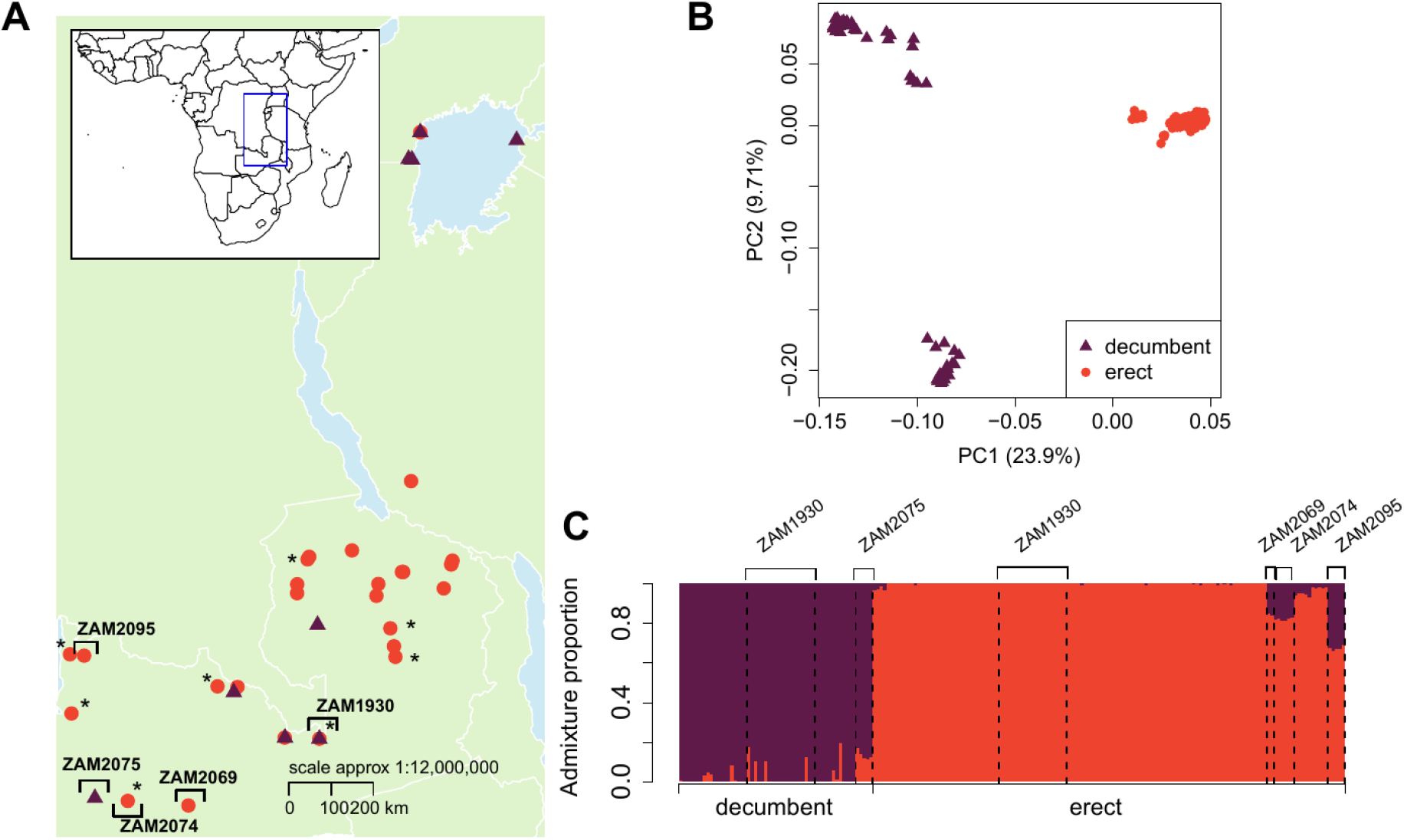
Population structure of *Alloteropsis angusta* in Zambia, Tanzania and Uganda. **A)** Geographic distribution of populations of *A. angusta* which were sampled and genotyped using RAD-sequencing. Red circles indicate erect populations, while decumbent populations are shown with purple triangles. Populations highlighted with an asterisk were sympatric with *A. semialata*. **B)** Principal component analysis of genetic variation among all RAD-sequenced *A. angusta* samples. **C)** Results of an admixture analysis at *K =* 2. Each vertical bar represents one individual. The major genetic groups are delimited at the bottom. Populations of interest are delimited by dashed vertical lines, with names at the top.

Pairwise comparisons of genetic differentiation (*F*_ST_) between populations reveal clear isolation-by-distance within both the decumbent (Mantel test: *ρ* = 0.84, *P* < 0.001) and erect (Mantel test: *ρ* = 0.43, *P* = 0.0019; Fig. 4) groups. However, the genetic divergence between populations representing distinct ecotypes did not increase with geographic distance (Mantel test: *ρ* = 0.026, *P* = 0.45; Fig. 4). This indicates ongoing gene flow between nearby populations within each ecotype, but amounts of gene flow that do not markedly increase between neighbouring erect and decumbent populations. Intermediate levels of genetic differentiation are present between pairs of erect and decumbent populations from Zambia and Tanzania (genome-wide average *F*_ST_: 0.399 – 0.568), which appears to result from many loci spread across the genome (Fig. S7).

**Figure 4.**
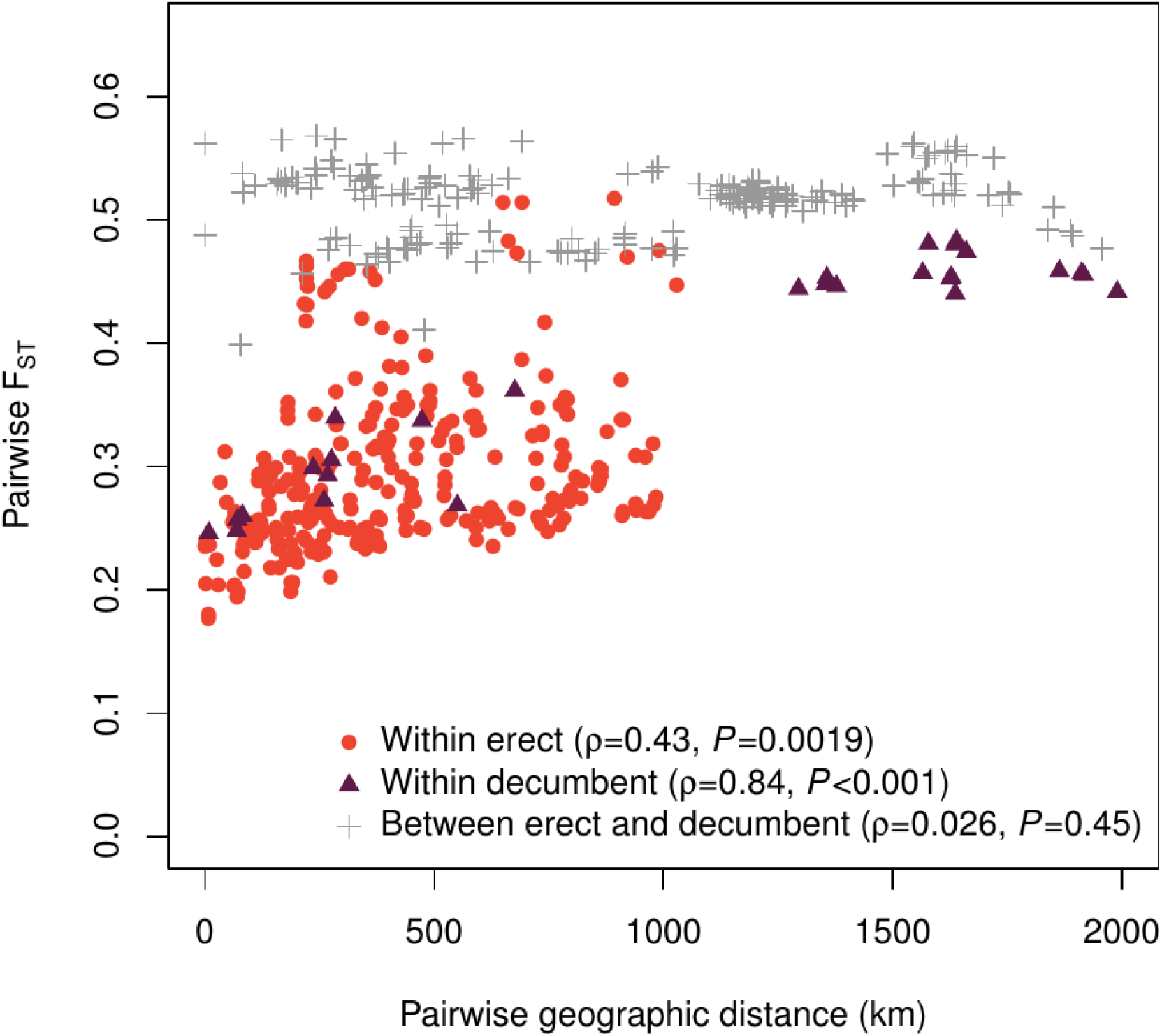
Patterns of isolation-by-distance within and between the two ecotypes of *Alloteropsis angusta*. Pairwise genetic differentiations (*F*_ST_) among all pairs of populations are plotted against geographic distances (km). Red circles indicate comparisons between erect populations, purple triangles indicate comparisons between decumbent populations, and grey crosses indicate comparisons between contrasting ecotypes. For each group, the Spearman correlation coefficient is indicated, with the *p-*value from a Mantel test.

### *Gene flow between the two ecotypes, and between* A. angusta *and* A. semialata

The ABBA-BABA tests revealed extensive gene flow between the erect and decumbent forms of *A. angusta* (Fig. 5). When the least often introgressed erect accession was put in the P1 position (TAN1; Fig. 3B), all tests with an erect in the P2 position and a decumbent in the P3 position were significant (Fig. 5A). Conversely, when the least often introgressed decumbent accession was put in the P1 position (UGA1; Fig. 5B), only some of the tests with a decumbent in the P2 position and an erect in the P3 position were significant (Fig. 5A). The highest *D*-statistics were observed between the erect ZAM2074-15 and the decumbent ZAM2075-04. Because other accessions from each group also showed elevated *D*-statistics with these two individuals, the exchanges were likely bidirectional. Estimates of admixture proportions using population-level data also support introgression between ecotypes in these two populations (Fig. 3C). In addition, the erect COD1 likely received genes from an unsampled decumbent lineage, explaining that it appears similarly introgressed by all decumbent individuals, and the lineages leading to the erect TAN1601-51 and the decumbent UGA4 likely exchanged genes (Fig. 5A). Other exchanges are difficult to pinpoint precisely, but significant comparisons involving two closely related decumbent (ZAM1930-17 and ZAM1950-10) and two or three of the closely-related erect individuals (ZAM1720-04, ZAM1933-01 and ZAM1930-JKO0102) suggest either exchanges among their common ancestors or repeated exchanges among each of them (Fig. 5A). Importantly, there is no strong evidence of exchanges specifically between the erect and decumbent individuals from the same location with very similar chloroplast genomes (ZAM1930-JKO0102 and ZAM1930-17; Fig. 5A).

**Figure 5.**
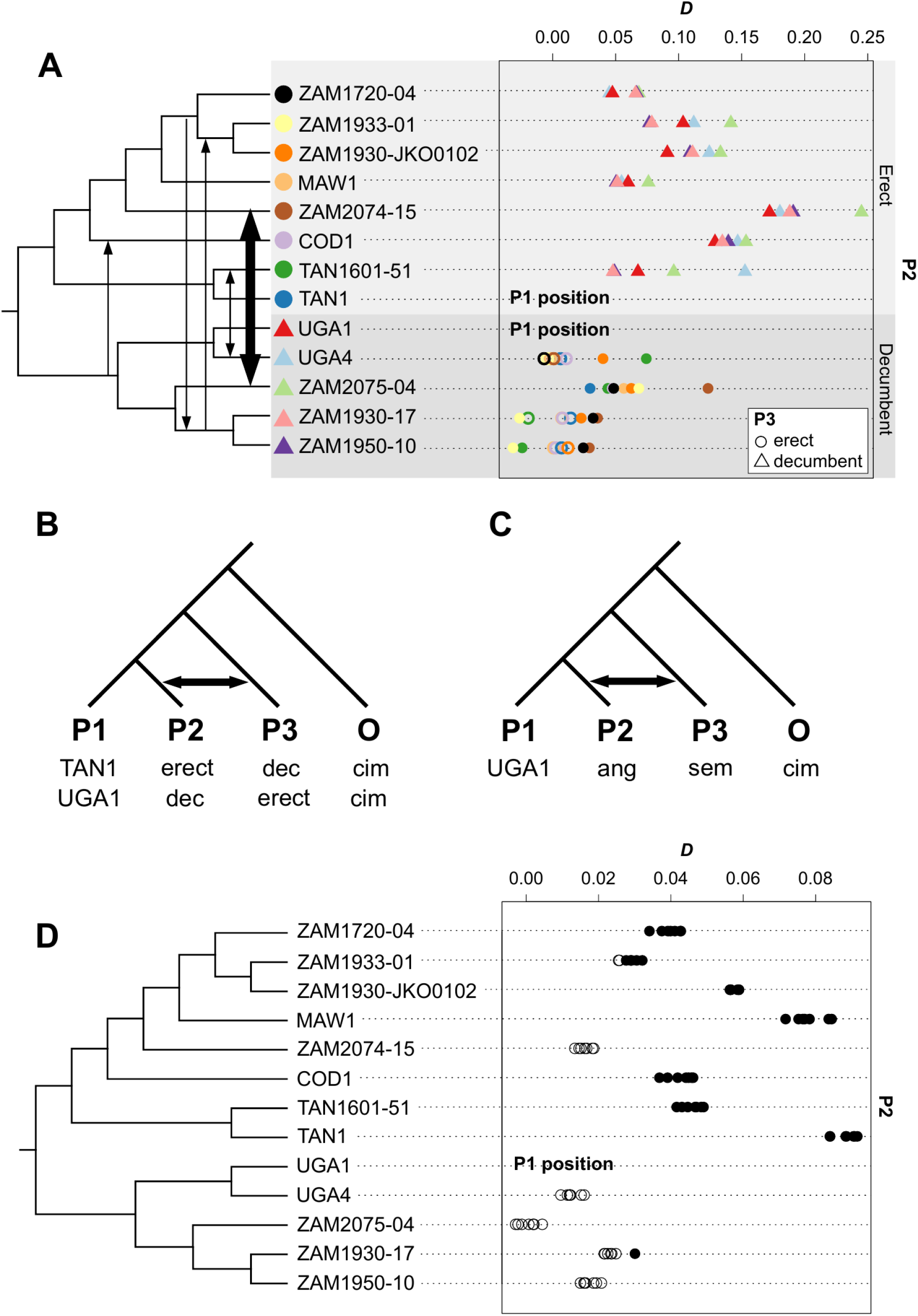
Evidence of introgression in *Alloteropsis angusta*. **A)** Distribution of *D*-statistics for each individual in the position P2 (ordered as in the multigene coalescence tree; Figure 2D), when the P1 position was occupied by the least often introgressed individual from the same ecotype and the P3 position was occupied by an individual from the other ecotype. The identity of the individual in the P3 position is indicated with symbols that match those in front of the phylogeny. Empty symbols show non-significant tests, filled symbols show significant tests. Arrows on the phylogenetic tree show the main genetic exchanges suggested by the *D*-statistics. **B)** Configuration used to detect introgression among the two ecotypes of *A. angusta*, with generic names of the positions (first line), the configuration focused on the erect individuals (second line), and the configuration focused on the decumbent individuals (third line; dec = decumbent, cim = *A. cimicina*). The arrow shows exchanges indicated by a positive *D*-statistic. **C)** Configuration used to detect introgression between *A. angusta* and *A. semialata*, using the same structure as in panel **B** (ang = *A. angusta*, sem = *A. semialata*). **D)** Distribution of *D*-statistics for each *A. angusta* individual in the P2 position (least often introgressed *A. angusta* individual in P1 position, *A. semialata* in P3 position; see structure in panel **C**). Individuals are arranged as in the multigene coalescence species tree (see Figure 2D). Empty circles indicate non-significant tests, with significant tests shown with filled circles.

ABBA-BABA tests were also conducted to test for introgression between *A. angusta* and its sister species *A. semialata*. When compared to the least often introgressed *A. angusta* from Uganda (UGA1; Fig. 5C), *D*-statistics revealed a significant excess of shared derived alleles between all *A. semialata* individuals and seven of the thirteen *A. angusta* (Fig. 5D). With the exception of ZAM2074-15, all erect individuals showed significant introgression from *A. semialata*, while none was detected in the decumbent individuals (except for one of the comparisons involving ZAM1930-17; Fig. 5D). These results indicate that genetic exchanges happened repeatedly between *A. semialata* and *A. angusta*, always involving the erect form of the latter (Fig. 5D), which presents similar ecological niches and growth habits as *A. semialata* (Fig. 1).

## Discussion

We analysed the genetic structure of two newly identified ecotypes of the grass *Alloteropsis angusta*; a decumbent ecotype restricted to wetlands and an erect ecotype growing in the miombo woodlands and grasslands of tropical Africa (Figs 1B, 1C, S1). The chloroplast genomes are shared among distinct ecotypes that are geographically close. By contrast, the nuclear genomes of the two ecotypes are deeply divergent, despite evidence of secondary gene flow between them. Population genomics of a densely sampled region confirms that the two ecotypes behave as two independent nuclear groups, with only a few admixture events. These patterns are likely the result of pollen-mediated hybridisation followed by movement of the hybrids into a different habitat, where selection favours the paternal genome. Our work therefore provides empirical evidence for a role of hybridisation in facilitating the dispersal of contrasted ecotypes across a large region of Africa.

### Selection maintains two contrasted ecotypes despite gene flow

Despite their geographic overlap, the two ecotypes of *A. angusta* are placed in strongly divergent groups in all analyses of nuclear genomes (Figures 2, 3 and 4). The ancestral state of the species is unknown, but coalescence analyses indicate a higher level of allele sorting at the base of the decumbent group (Fig. 2D). This might indicate that the decumbent lineage emerged from within a large erect clade and then underwent an initial bottleneck. The erect and bulbous habit is shared with *A. semialata*, which could support the presence of these characters in their common ancestor. Independently of the direction of the transition, the emergence of a new growth habit has enabled the colonisation of a new habitat, as the two ecotypes are consistently associated with distinct habitats (Figs S1, S2). The population-level analyses identified cases of admixture between some of the erect and decumbent populations (Fig. 3C), and the ABBA-BABA tests revealed multiple cases of introgression among the two ecotypes (Fig. 5). These results demonstrate that the two groups have recurrently interbred. The marked genetic structure must therefore result from strong habit-specific selection. Our scans identify peaks of differentiation distributed throughout the genome (Fig. S7), suggesting that selection has acted on multiple loci. These regions might be involved in dimensions of the phenotype other than the growth habit, as the contrasted habitats will also differ in their light, water, and nutrient availability. Our analyses therefore suggest selection maintained two different life strategies across a vast area despite gene flow.

### Hybridisation allows co-dispersal of ecotypes

Unlike the nuclear genome, chloroplast variants are sorted geographically, independently of the ecotype (Fig. 2B). In particular, decumbent and erect individuals occurring across a miombo-wetland boundary in Zambia share almost identical chloroplasts (population ZAM1930; Figs 2B, S2). Because of their smaller effective population size, haploid chloroplast genomes are more easily introgressed (Martinsen *et al*., 2001). Frequent chloroplast sharing might therefore result from hybridisation among geographically close populations, a process usually referred to as cytoplasmic capture and reported in multiple species of trees (e.g. Petit et al., 1997; Thomson *et al*., 2015; Gryta *et al*., 2017). However, chloroplast-nuclear discrepancies might be better explained by dispersal dynamics (Petit *et al*., 2003). In most species, the maternally-inherited chloroplast genomes are transported solely by seeds, while nuclear genomes are transported by both seeds and pollen. As a grass, *A. angusta* is anemophile and its seeds do not present obvious dispersal mechanisms. The ecological specificity of the two ecotypes, combined with a patchy distribution of each environment at the studied scale, means that suitable habitats for either ecotype might be separated by large distances. In wind-dispersed trees, pollen travels further than seeds (Heuertz *et al*., 2003; Bittencourt & Sebbenn, 2007), and pollen-mediated gene flow can occur over large distances in grasses (Busi *et al*., 2008). In such instances, frequent hybridisation should lead to mixing of the nuclear genomes (Wallace *et al*., 2011; Aguilée *et al*., 2016). We observe the opposite here, and suggest that episodic long-distance pollen-mediated hybridisation, followed by strong habitat-specific selection, allows the dispersal of one ecotype via the seeds from the other ecotype, as suggested in some trees (Potts & Reid, 1988; Petit *et al*., 2003). This dispersal mechanism provides a shortcut to the slower process of colonisation by seed. Previous examples concern morphotypes that grow sympatrically in the same habitat types, but our analyses of *A. angusta* show that a similar process enhances the dispersal capabilities of ecotypes otherwise separated by habitat-specific selection.

Following long-distance pollination of one established population by the other ecotype, hybrids would bear the local-type chloroplast with a mixed nuclear genome. Their paternal alleles adapted to the other habitat might allow its colonisation. Following either crosses among hybrids or further long-distance pollination by the other ecotype, selection would repeatedly favour the alleles corresponding to the new habitat. Over time, this process will result in the combination of nuclear genomes mostly matching the new habitat with chloroplast genomes originating from the other habitat. An ecotype can therefore effectively colonise a suitable habitat by hijacking the seeds from the contrasting ecotype, leaving behind only small traces of hybridisation, as detected in some populations by the introgression tests (Fig. 5). In the population with both decumbent and erect individuals (ZAM1930; Fig. S2), admixture was suggested for a few decumbent individuals, but none of the erect individuals (Fig. 3C). In addition, the two individuals with whole-genome sequencing did not show clear evidence of direct exchanges among them (Fig. 5A). In this case, the hybridisation that allowed the exchange of chloroplast genomes left very little traces in the nuclear genome, showing that the alleles from the other type of habitat can be rapidly purged.

The mixture of erect and decumbent types in the chloroplast phylogenetic tree (Fig. 2B) suggests that such co-dispersal, potentially coupled with post-dispersal introgression, occurred recurrently. The decumbent type is associated with rivers that massively increase in volume during the rainy season, likely providing ample opportunities for unidirectional long-distance seed transport. Distant decumbent populations might then provide a gateway for colonisation of distant habitats by the erect form, contributing to the overall spread of *A. angusta*. Conversely, hybridisation with the erect form growing in woodlands and grasslands would provide access to unconnected river bodies or suitable sites located upstream. We conclude that hybridisation among ecotypes inhabiting markedly different habitats increases the dispersal potential of *A. angusta*.

### *Interspecific exchanges increase diversity within* A. angusta

Besides the two *A. angusta* ecotypes, the conspecific *A. semialata* is also frequent in the region and in several instances was found growing mixed with erect *A. angusta* (Table S1). Our clustering analyses provide evidence of admixture between *A. semialata* and *A. angusta* in two individuals, always involving the erect ecotype of the latter (Fig. 3). In addition, ABBA-BABA tests indicate that the erect individuals of *A. angusta* are significantly more introgressed than their decumbent conspecifics (Fig. 5). The erect form of *A. angusta* and *A. semialata* co-occur frequently and share the same habitats, which would increase opportunities for hybridisation. In addition, the similarity of growth forms and habitats might have favoured the sharing of adaptive alleles via selective introgression. Some individuals of *A. angusta* are very similar to *A. semialata* (Fig. 1), with large deep bulbs that are not usually found in *A. angusta*. Genes responsible for such traits might have crossed the species boundaries, as reported for elements of the C_4_ photosynthetic pathway (Dunning *et al*., 2017). The two species therefore form a species complex, with recurrent genetic exchanges between the two ecotypes of *A. angusta* (and the different subgroups of *A. semialata*; Bianconi *et al*., 2020; Olofsson *et al*., 2021), but also episodic exchanges among *A. semialata* and *A. angusta*. The impact of these exchanges remains speculative, but they might have contributed to the functional and ecological diversity of the group.

## Conclusions

In this work, we show that the grass *A. angusta* exists across tropical Africa as two contrasted ecotypes that correspond to distinct genetic groups. The erect form, previously mistaken for the conspecific *A. semialata*, is widespread in miombo woodlands, while the decumbent form occurs in wetlands bordering lakes and rivers. Despite deep nuclear divergence, we find evidence of genetic exchanges and different morphs from a given geographical region share chloroplast genomes. These patterns indicate recurrent hybridisation followed by selection that sorts most of the genomes by habitat. These hybridisation events offer opportunities for dispersal to distant locations by effectively hijacking the seeds from the other ecotype. We conclude that hybridisation, coupled with strong selection, can boost plant dispersal without erasing the associations between genomes and environments. A similar mechanism has been previously proposed among tree morphotypes, and we offer here empirical evidence of co-dispersal among grasses adapted to contrasting habitats.

## Acknowledgements

This work was funded by the European Research Council (grant ERC-2014-STG-638333) and the Royal Society (grant RGF\EA\181050), and has benefited from “Investissements d’Avenir” grants managed by the Agence Nationale de la Recherche (CEBA, ref. ANR-10-LABX-25-01 and TULIP, ref. ANR-10-LABX-41). P.A.C. is funded by a Royal Society University Research Fellowship (grant URF\R\180022).

## Author contributions

EVC, MSS, JKO and PAC designed the study. EVC, JKO, FN, MEB and PAC performed field work. EVC, MSS, JKO, GS, SM, and GB produced the genomic data. EVC analysed the data, with the help of MSS, JKO, MEB, and PAC. EVC and PAC wrote the paper, with the help of all authors.

## Data Availability

Sequence data have been deposited in the NCBI Sequence Read Archive with the project number PRJNA715711. Scripts for analysis and data processing available at: https://github.com/evcurran/Angusta-Pop-Genomics

## References

Abbott R, Albach D, Ansell S, Arntzen JW, Baird SJE, Bierne N, Bougham J, Brelsford A, Buerkle CA, Buggs R et al. 2013. Hybridization and speciation. Journal of Evolutionary Biology 26: 229–246.

Aguilée R, Raoul G, Rousset F, Ronce O. 2016. Pollen dispersal slows geographical range shift and accelerates ecological niche shift under climate change. Proceedings of the National Academy of Sciences of the United States of America, 113: E5741–E5748.

Arnold BJ, Lahner B, DaCosta JM, Weisman CM, Hollister JD, Salt DE, Bomblies K, Yant L. 2016. Borrowed alleles and convergence in serpentine adaptation. Proceedings of the National Academy of Sciences of the United States of America, 113: 8320–8325.

Barton NH, Hewitt GM. 1985. Analysis of hybrid zones. Annual Review of Ecology and Systematics, 16: 113–148.

Bhatia G, Patterson N, Sankararaman S, Price AL. 2013. Estimating and interpreting FST: The impact of rare variants. Genome Research 23: 1514–1521.

Bianconi ME, Dunning LT, Curran EV, Hidalgo O, Powell RF, Mian S, Leitch IJ, Lundgren MR, Manzi S, Vorontsova MS, et al. Contrasted histories of organelle and nuclear genomes underlying physiological diversification in a grass species. Proceedings of the Royal Society B: Biological Sciences 287: 20201960. https://doi.org/10.1098/rspb.2020.1960.

Bittencourt JVM, Sebbenn AM. 2007. Patterns of pollen and seed dispersal in a small, fragmented population of the wind-pollinated tree Araucaria angustifolia in southern Brazil. Heredity 99: 580–591.

Bolger AM, Lohse M, Usadel B. 2014. Trimmomatic: A flexible trimmer for Illumina sequence data. Bioinformatics 30: 2114–2120.

Broad Institute. 2019. Picard Toolkit. GitHub Repository. http://broadinstitute.github.io/picard/;BroadInstitute

Busi R, Yu Q, Barrett-Lennard R, Powles S. 2008. Long distance pollen-mediated flow of herbicide resistance genes in Lolium rigidum. Theoretical and Applied Genetics 117: 1281–1290.

Capella-Gutiérrez S, Silla-Martínez JM, Gabaldón T. 2009. trimAl: A tool for automated alignment trimming in large-scale phylogenetic analyses. Bioinformatics 25: 1972– 1973.

Catchen J, Hohenlohe PA, Bassham S, Amores A, Cresko WA. 2013. Stacks: An analysis tool set for population genomics. Molecular Ecology 22: 3124–3140.

Clayton WD, Renvoize SA. 1982. Flora of tropical East Africa. Gramineae. Part 3. AA Balkema, Rotterdam.

Drummond AJ, Rambaut A. 2007) BEAST: Bayesian evolutionary analysis by sampling trees. BMC Evolutionary Biology 7: 214.

Dunning LT, Lundgren MR, Moreno-Villena JJ, Namaganda M, Edwards EJ, Nosil P, Osborne CP, Christin PA. 2017. Introgression and repeated co-option facilitated the recurrent emergence of C4 photosynthesis among close relatives. Evolution 71: 1541– 1555.

Dunning LT, Olofsson JK, Parisod C, Choudhury RR, Moreno-Villena JJ, Yang Y, Dionora J, Quick PW, Park M, Bennetzen JL et al. 2019. Lateral transfers of large DNA fragments spread functional genes among grasses. Proceedings of the National Academy of Sciences of the United States of America 116: 4416–4425.

Durand EY, Patterson N, Reich D, Slatkin M. 2011. Testing for ancient admixture between closely related populations. Molecular Biology and Evolution 28: 2239–2252.

Ellstrand NC. 1992. Gene flow by pollen: Implications for plant conservation genetics. Oikos 63: 77.

Ellstrand NC. 2014. Is gene flow the most important evolutionary force in plants? American Journal of Botany 101: 737–753.

Evanno G, Regnaut S, Goudet J. 2005. Detecting the number of clusters of individuals using the software STRUCTURE: A simulation study. Molecular Ecology 14: 2611–2620.

Galindo J, Martínez-Fernández M, Rodríguez-Ramilo ST, Rolán-Alvarez E. 2013. The role of local ecology during hybridization at the initial stages of ecological speciation in a marine snail. Journal of Evolutionary Biology 26: 1472–1487.

Green RE, Krause J, Briggs AW, Maricic T, Stenzel U, Kircher M, Patterson N, Li H, Zhai W, Fritz MHS, et al. 2010. A draft sequence of the neandertal genome. Science 328: 710–722.

Grass Phylogeny Working Group. 2001. Phylogeny and subfamilial classification of the grasses (Poaceae). Annals of the Missouri Botanical Garden 88: 373–457.

Gryta H, Van de Paer C, Manzi S, Holota H, Roy M, Besnard G. 2017. Genome skimming and plastid microsatellite profiling of alder trees (Alnus spp., Betulaceae): phylogenetic and phylogeographical prospects. Tree Genetics and Genomes 13: 118.

Hall RJ. 2016. Hybridization helps colonizers become conquerors. Proceedings of the National Academy of Sciences of the United States of America 113: 9963–9964.

Heuertz M, Vekemans X, Hausman JF, Palada M, Hardy OJ. 2003. Estimating seed vs. pollen dispersal from spatial genetic structure in the common ash. Molecular Ecology 12: 2483–2495.

Hudson RR, Slatkin M, Maddison WP. 1992. Estimation of levels of gene flow from DNA sequence data. Genetics 132: 583–589.

Ibrahim DG, Gilbert ME, Ripley BS, Osborne CP. 2008. Seasonal differences in photosynthesis between the C3 and C4 subspecies of Alloteropsis semialata are offset by frost and drought. Plant, Cell & Environment 31: 1038–1050. https://doi.org/10.1111/j.1365-3040.2008.01815.x

Kellogg EA. 2015. Flowering Plants. Monocots: Poaceae. Heidelberg: Springer. p. 416.

Kopelman NM, Mayzel J, Jakobsson M, Rosenberg NA, Mayrose I. 2015. Clumpak: A program for identifying clustering modes and packaging population structure inferences across K. Molecular Ecology Resources 15: 1179–1191.

Korneliussen TS, Albrechtsen A, Nielsen R. 2014. ANGSD: Analysis of next generation sequencing data. BMC Bioinformatics 15: 1–13.

Langmead B, Salzberg SL. 2012. Fast gapped-read alignment with Bowtie 2. Nature Methods 9: 357–359.

Li H. 2011. A statistical framework for SNP calling, mutation discovery, association mapping and population genetical parameter estimation from sequencing data. Bioinformatics, 27: 2987–2993.

Linder HP, Lehmann CER, Archibald S, Osborne CP, Richardson DM. 2018. Global grass (Poaceae) success underpinned by traits facilitating colonization, persistence and habitat transformation. Biological Reviews 93: 1125–1144.

Lu G, Basley DJ, Bernatchez L. 2001. Contrasting patterns of mitochondrial DNA and microsatellite introgressive hybridization between lineages of lake whitefish (Coregonus clupeaformis); relevance for speciation. Molecular Ecology 10: 965–985.

Lundgren MR, Besnard G, Ripley BS, Lehmann CER, Chatelet DS, Kynast RG, Namaganda M, Vorontsova MS, Hall RC, Elia J, et al. 2015. Photosynthetic innovation broadens the niche within a single species. Ecology Letters 18: 1021–1029.

Martinsen GD, Whitham TG, Turek RJ, Keim P. 2001. Hybrid populations selectively filter gene introgression between species. Evolution 55: 1325–1335.

Meisner J, Albrechtsen A. 2018. Inferring population structure and admixture proportions in low-depth NGS data. Genetics 210: 719–731.

Nychka D, Furrer R, Paige J, Sain S. 2017. “fields: Tools for spatial data.” doi: 10.5065/D6W957CT

O’Connell J, Schulz-Trieglaff O, Carlson E, Hims MM, Gormley NA, Cox AJ. 2015. NxTrim: Optimized trimming of Illumina mate pair reads. Bioinformatics 31: 2035– 2037.

Olofsson JK, Bianconi M, Besnard G, Dunning LT, Lundgren MR, Holota H, Vorontsova MS, Hidalgo O, Leitch IJ, Nosil P, et al. 2016. Genome biogeography reveals the intraspecific spread of adaptive mutations for a complex trait. Molecular Ecology 25: 6107–6123.

Olofsson JK, Dunning LT, Lundgren MR, Barton HJ, Thompson J, Cuff N, Anyarathne M, Yakandawala D, Sotelo G, Zeng K, et al. 2019. Population-specific selection on standing variation generated by lateral gene transfers in a grass. Current Biology 29: 3921-3927.e5.

Olofsson JK, Curran EV, Nyirenda F, Bianconi ME, Dunning LT, Milenkovic V, Sotelo G, Hidalgo O, Powell RF, Lundgren MR, et al. 2021. Low dispersal and ploidy differences in a grass maintain photosynthetic diversity despite gene flow and habitat overlap. Molecular Ecology https://doi.org/10.1111/mec.15871

Patel RK, Jain M. 2012. NGS QC toolkit: A toolkit for quality control of next generation sequencing data. PLoS ONE 7: e30619.

Perreta M, Ramos J, Tivano JC, Vegetti A. 2011. Descriptive characters of growth form in Poaceae - An overview. Flora: Morphology, Distribution, Functional Ecology of Plants 206: 283–293.

Petit RJ, Bodénès C, Ducousso A, Roussel G, Kremer A. 2003. Hybridization as a mechanism of invasion in oaks. New Phytologist 161: 151–164.

Petit RJ, Pineau E, Demesure B, Bacilieri R, Ducousso A, Kremer A. 1997. Chloroplast DNA footprints of postglacial recolonization by oaks. Proceedings of the National Academy of Sciences of the United States of America 94: 9996–10001.

Potts BM, Reid JB. 1988. Hybridization as a dispersal mechanism. Evolution 42: 1245–1255.

R Core Team. 2019. R: A language and environment for statistical computing. R Foundation for Statistical Computing, Vienna, Austria. URL https://www.R-project.org/

Rambaut A, Suchard MA, Xie W, Drummond AJ. 2013. Tracer v1.6. Available from: http://tree.bio.ed.ac.uk/software/tracer/

Schluter D. 2001. Ecology and the origin of species. Trends in Ecology & Evolution 16: 372– 380.

Schneider CA, Rasband WS, Eliceiri KW. 2012. NIH Image to ImageJ: 25 years of image analysis. Nature Methods 9: 671–675.

Seehausen O, Butlin RK, Keller I, Wagner CE, Boughman JW, Hohenlohe PA, Peichel CL, Saetre GP, Bank C, Brännström Å, et al. 2014. Genomics and the origin of species. Nature Reviews Genetics 15: 176–192.

Skotte L, Korneliussen TS, Albrechtsen A. 2013. Estimating individual admixture proportions from next generation sequencing data. Genetics 195: 693–702.

Song Y, Endepols S, Klemann N, Richter D, Matuschka FR, Shih CH, Nachman MW, Kohn MH. 2011. Adaptive introgression of anticoagulant rodent poison resistance by hybridization between old world mice. Current Biology 21: 1296–1301.

Soria-Carrasco V, Gompert Z, Comeault AA, Farkas TE, Parchman TL, Johnston JS, Buerkle CA, Feder JL, Bast J, Schwander T, et al. 2014. Stick insect genomes reveal natural selection’s role in parallel speciation. Science 344: 738–742.

Stamatakis A. 2014. RAxML version 8: A tool for phylogenetic analysis and post-analysis of large phylogenies. Bioinformatics 30: 1312–1313.

Stankowski S, Sobel JM, Streisfeld MA. 2017. Geographic cline analysis as a tool for studying genome-wide variation: a case study of pollinator-mediated divergence in a monkeyflower. Molecular Ecology 26: 107–122.

Stapf O. 1919. In Daniel, O. 1919. Flora of Tropical Africa. Crown Agents for Overseas Governments and Administrations, London. pp. 482–486.

Stent SM, Rattray JM. 1933. The grasses of southern Rhodesia. Proceedings and Transactions of the Rhodesia Scientific Association. Bulawayo 32: 21–22.

Taylor EB, Boughman JW, Groenenboom M, Sniatynski M, Schluter D, Gow JL. 2006. Speciation in reverse: Morphological and genetic evidence of the collapse of a three-spined stickleback (Gasterosteus aculeatus) species pair. Molecular Ecology 15: 343– 355.

The Heliconius Genome Consortium. 2012. Butterfly genome reveals promiscuous exchange of mimicry adaptations among species. Nature 487: 94–98.

Thomson AM, Dick CW, Pascoini AL, Dayanandan S. 2015. Despite introgressive hybridization, North American birches (Betula spp.) maintain strong differentiation at nuclear microsatellite loci. Tree Genetics and Genomes, 11: 101.

Todesco M, Pascual MA, Owens GL, Ostevik KL, Moyers BT, Hübner S, Heredia SM, Hahn MA, Caseys C, Bock DG, Rieseberg LH. 2016. Hybridization and extinction. Evolutionary Applications 9: 892–908.

Wallace LE, Culley TM, Weller SG, Sakai AK, Kuenzi A, Roy T, Wagner WL, Nepokroeff M. 2011. Asymmetrical gene flow in a hybrid zone of Hawaiian Schiedea (Caryophyllaceae) species with contrasting mating systems. PLoS ONE 6: 1–12.

Zhang C, Rabiee M, Sayyari E, Mirarab S. 2018. ASTRAL-III: Polynomial time species tree reconstruction from partially resolved gene trees. BMC Bioinformatics 19: 15–30.

