## Supplementary Material for "Hybridisation boosts dispersal of two contrasted ecotypes in a grass species"

This Supplementary Material contains one table and seven figures:

Table S1. Sample information (provided as a spreadsheet)

Figure S1. Photographs of individuals of each ecotype in the field

Figure S2. Distribution of individuals in population ZAM1930

Figure S3. Phylogenetic trees of *Alloteropsis*

Figure S4. Isolation by distance on plastid genomes

Figure S5. Coalescence species tree of *Alloteropsis*

Figure S6. Optimal number of genetic clusters in the dataset

Figure S7: Genomic landscape of differentiation between *Alloteropsis angusta* ecotypes

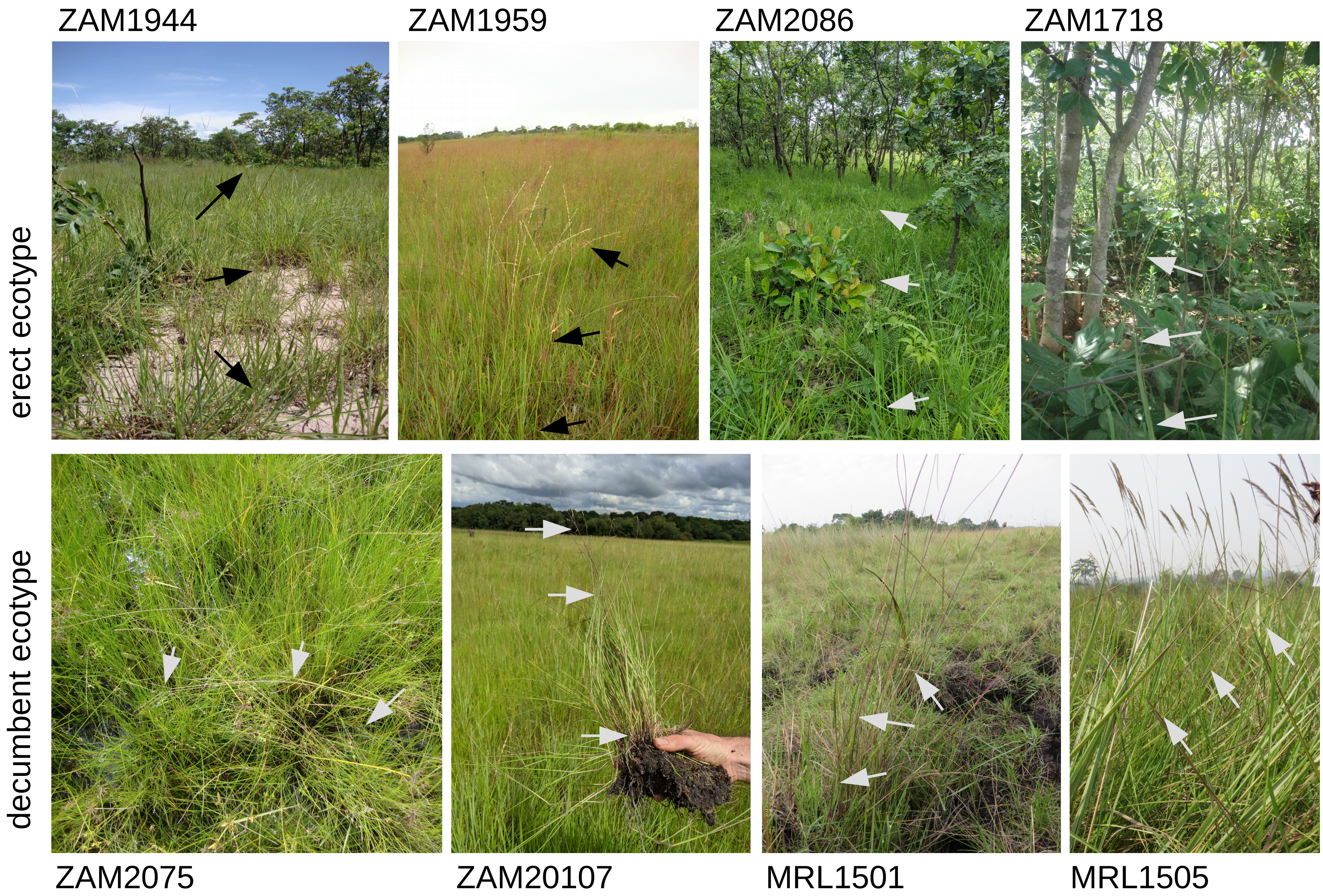


**Figure S1. Photographs of individuals of each ecotype in the field.** The top row of photographs shows individuals of the erect ecotype of *Alloteropsis angusta*, indicated by arrows. Population ZAM1944 occurred in a grassy opening in miombo woodlands, population ZAM1959 occurred in a grassland and populations ZAM2086 and ZAM1718 were both miombo woodlands. The bottom row of photographs shows individuals of the decumbent ecotype of *A. angusta*. ZAM2075 and ZAM20107 occurred in river wetlands and populations MRL1501 and MRL1505 grew among tall grasses.

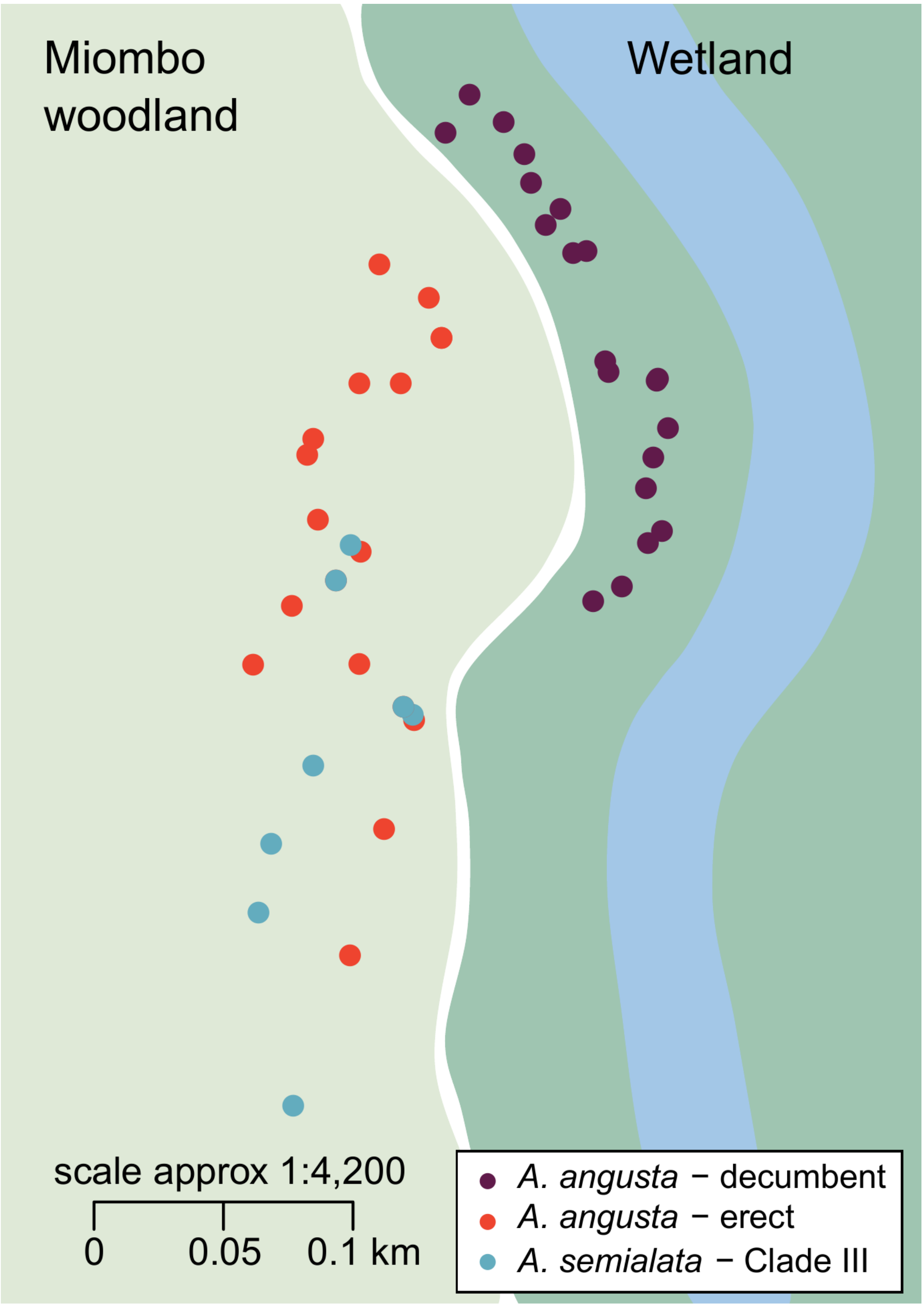


**Figure S2. Distribution of individuals in population ZAM1930.**
The position of individual plants is shown with circles, coloured per type. The different habitats are shown, with the river in blue, its associated wetland in green and the adjacent miombo woodland in light green.

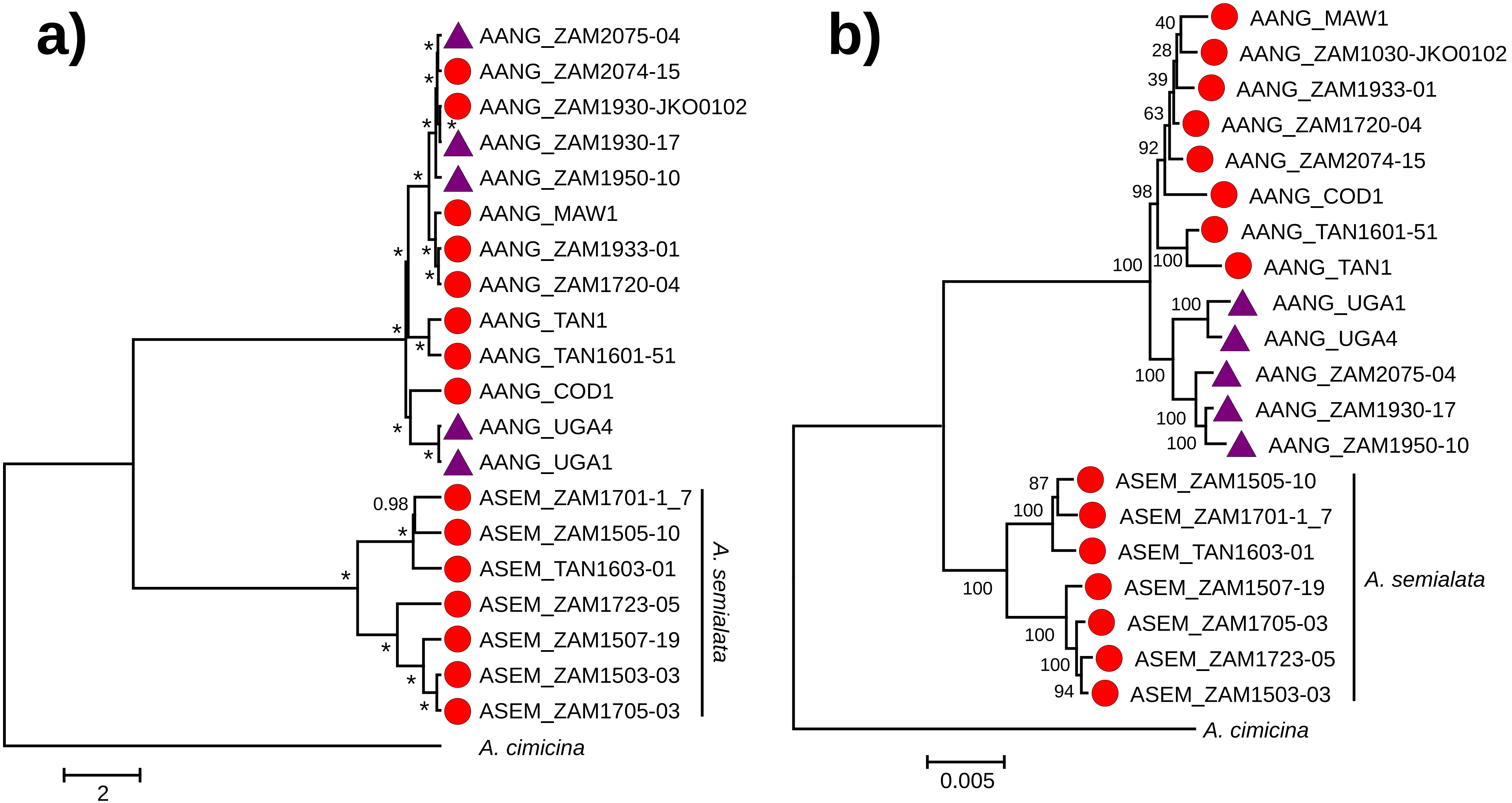


**Figure S3. Phylogenetic trees of *Alloteropsis*. a)** A time-calibrated tree was inferred from chloroplast genomes, with branch lengths in million years. Bayesian support values are shown near nodes; asterisks indicate nodes with a support of 1.0. **b)** A phylogram was inferred from nuclear variants, with branch lengths in expected substitutions per site. Bootstrap support values are indicated near nodes. For both panels, individuals growing erect from a bulb are shown with a red circle, while purple triangles show decumbent individuals.

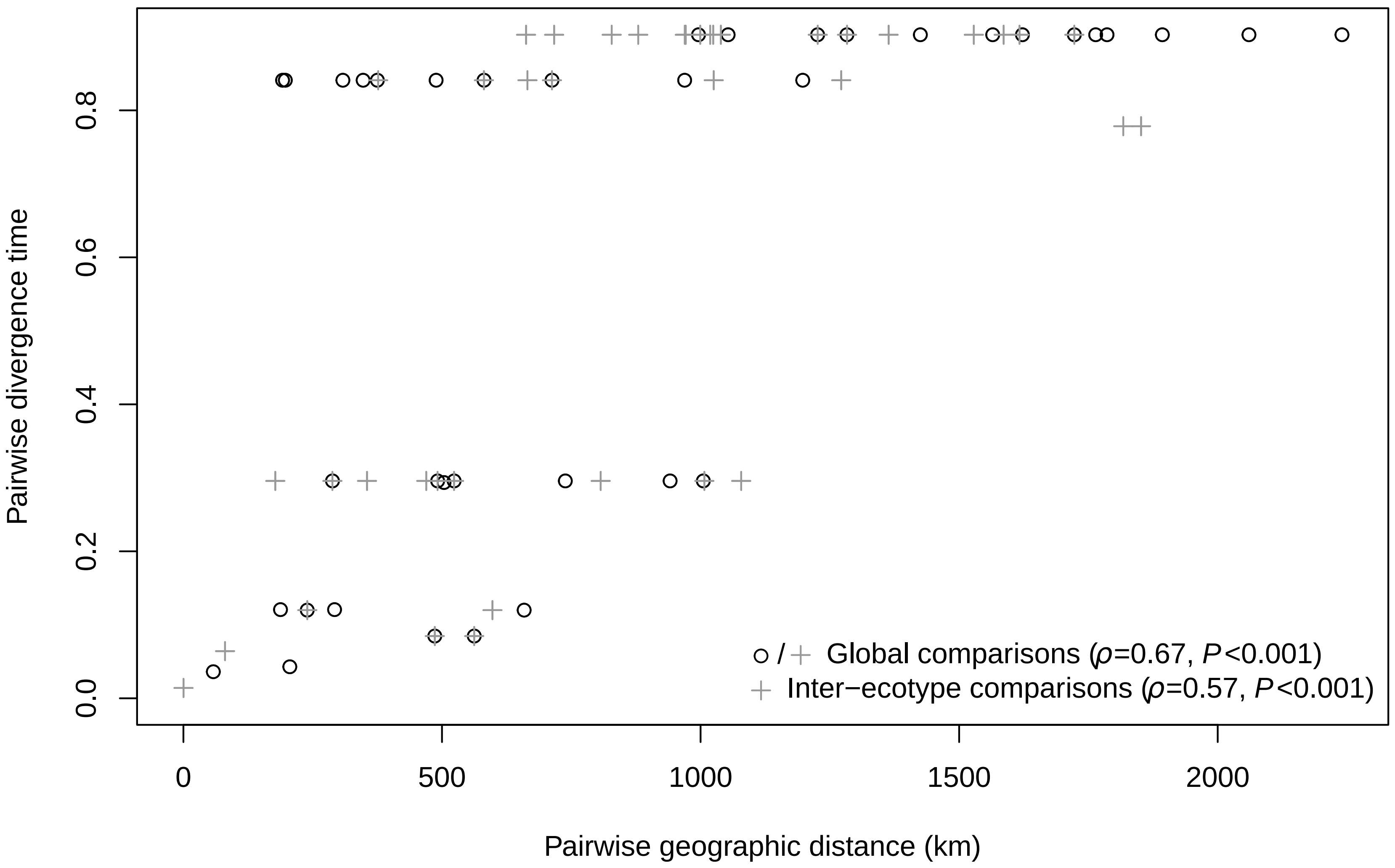


**Figure S4. Isolation by distance on plastid genomes.** Divergence times among all pairs of plastomes are plotted against geographic distances (km). Open circles indicate within-ecotype comparisons and grey crosses indicate between-ecotype comparisons. The Spearman correlation coefficient and the p-value from a Mantel test are indicate for both the global analysis between all populations and for the inter-ecotype comparisons.

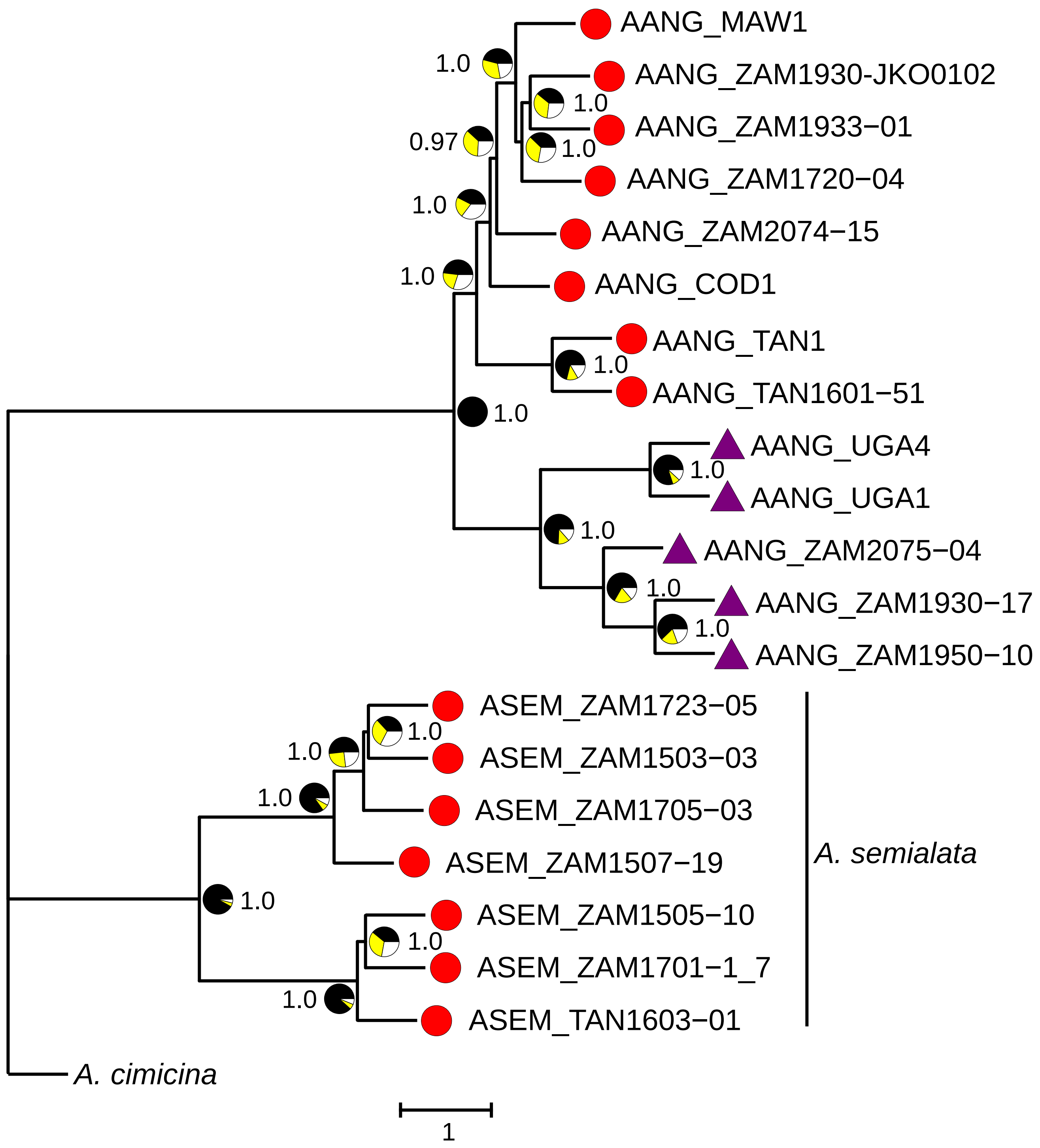


**Figure S5. Coalescence species tree of *Alloteropsis*.** The relationships were inferred from nuclear markers. Support values are indicated near nodes, with pie charts showing the proportion of quartets supporting the main topology (in black) and the two alternatives. Branch lengths are given in coalescence units. Terminal branches are given an arbitrary length. Individuals growing erect from a bulb are shown with a red circle, while purple triangles show decumbent individuals.

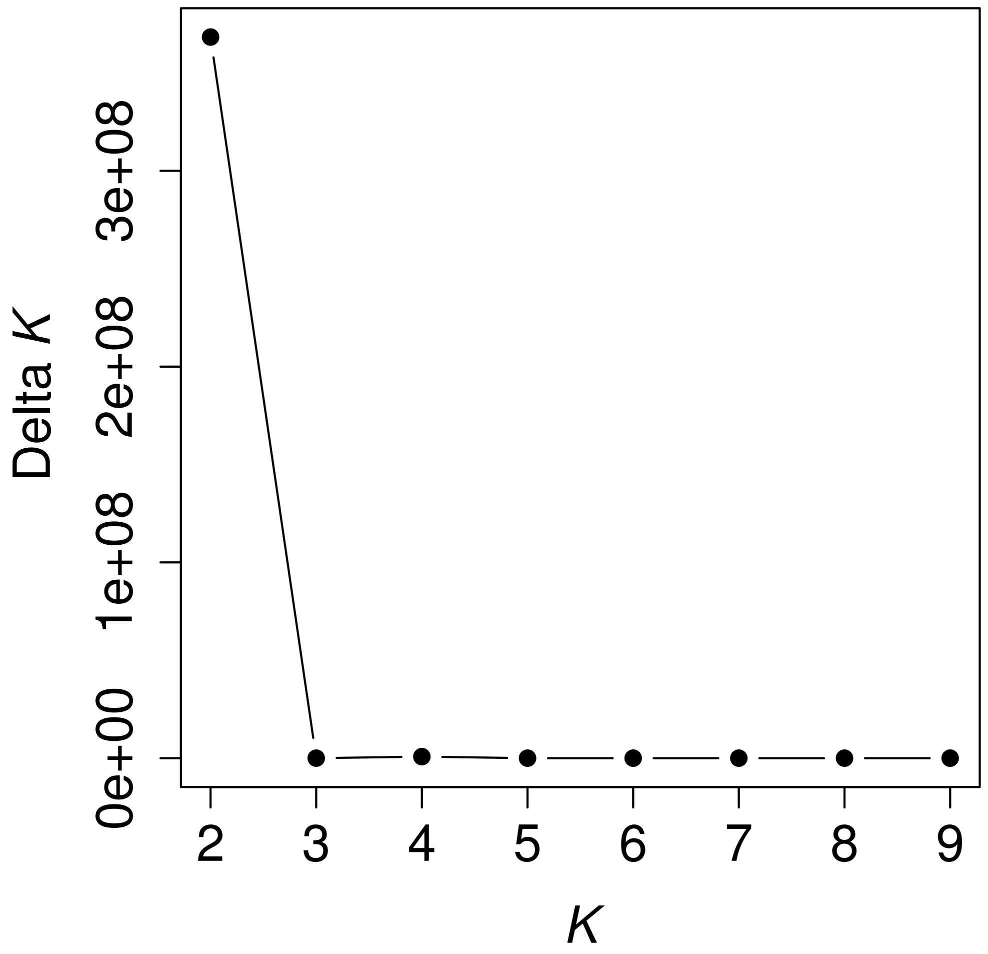


**Figure S6. Optimal number of genetic clusters in the dataset.** The delta-*K* method was used to determine the best value of *K* to describe population structure in the admixture analysis. The highest value of delta-*K* indicates the optimal value of *K*.

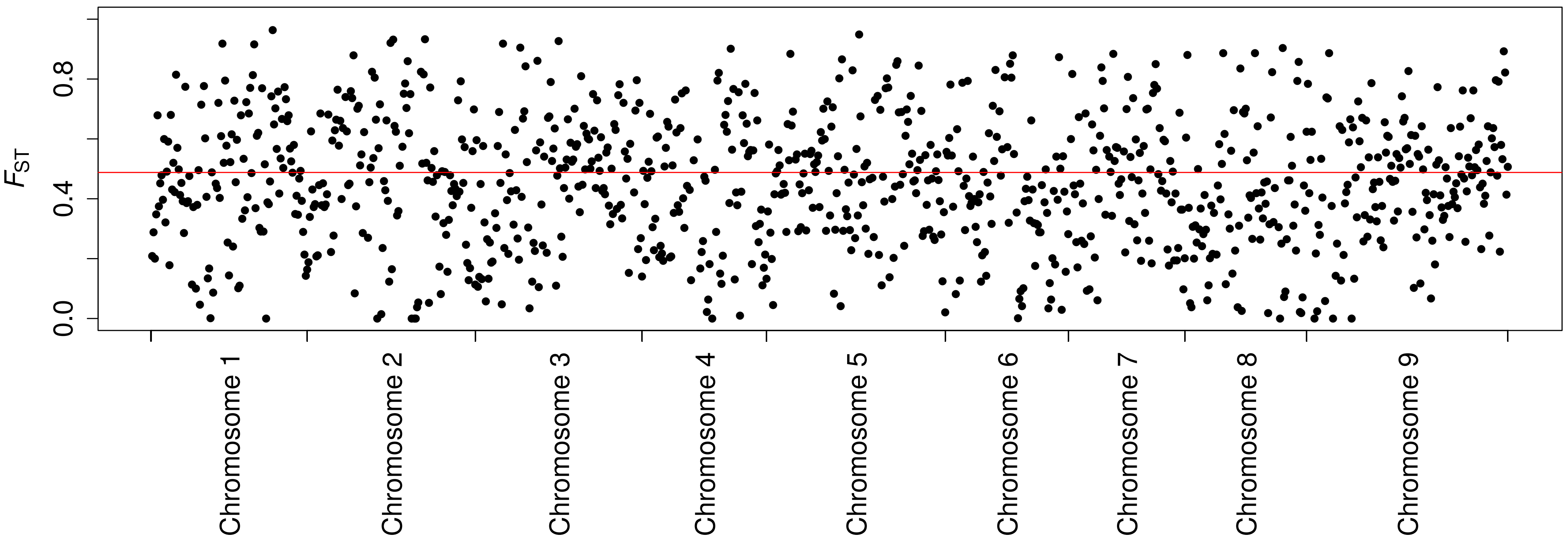


**Figure S7. Genomic landscape of differentiation between *Alloteropsis angusta* ecotypes.** The average $F_{ST}$ for 500kb windows is plotted along the genome, describing genetic differentiation between erect and decumbent ecotypes from the population ZAM19-30. The red horizontal line indicates the genome-wide average $F_{ST}$ of 0.488.
